# Splotch: Robust estimation of aligned spatial temporal gene expression data

**DOI:** 10.1101/757096

**Authors:** Tarmo Äijö, Silas Maniatis, Sanja Vickovic, Kristy Kang, Miguel Cuevas, Catherine Braine, Hemali Phatnani, Joakim Lundeberg, Richard Bonneau

**Author notes:** These authors contributed equally to this work (Tarmo Äijö, Silas Maniatis, Sanja Vickovic).

## Abstract

Spatial genomics technologies enable new approaches to study how cells interact and function in intact multicellular environments but present a host of technical and computational challenges. Here we describe Splotch, a novel computational framework for the analysis of spatially resolved transcriptomics data. Splotch aligns transcriptomics data from multiple tissue sections and timepoints to generate improved posterior estimates of gene expression. We demonstrate alignment of a large corpus of single-cell RNA-seq data into an automatically generated spatial-temporal coordinate and study optimal design for spatial transcriptomics experiments.

## Main

Unbiased spatial maps of gene expression are important resources for understanding diseases in intact tissue context^1–6^. Previous genome-wide studies have demonstrated the value of combining appropriate experimental designs with statistical and computational methods tailored to new genomic technologies^7,8^, and the study of spatial gene expression is no exception to this trend^9^. Here, we describe a method, Splotch, that combines integrative and spatiotemporal generative modeling (**Supplementary Fig. 1**) matched to the spatial transcriptomics (ST) technology^10,11^. Splotch enables interrogation of spatiotemporal genomics data by simultaneously using spatial, temporal and experimental (e.g. genotype) coordinates to improve probabilistic inference of spatially resolved gene expression quantities (**Fig. 1a**; *Methods*).

**Figure 1.**
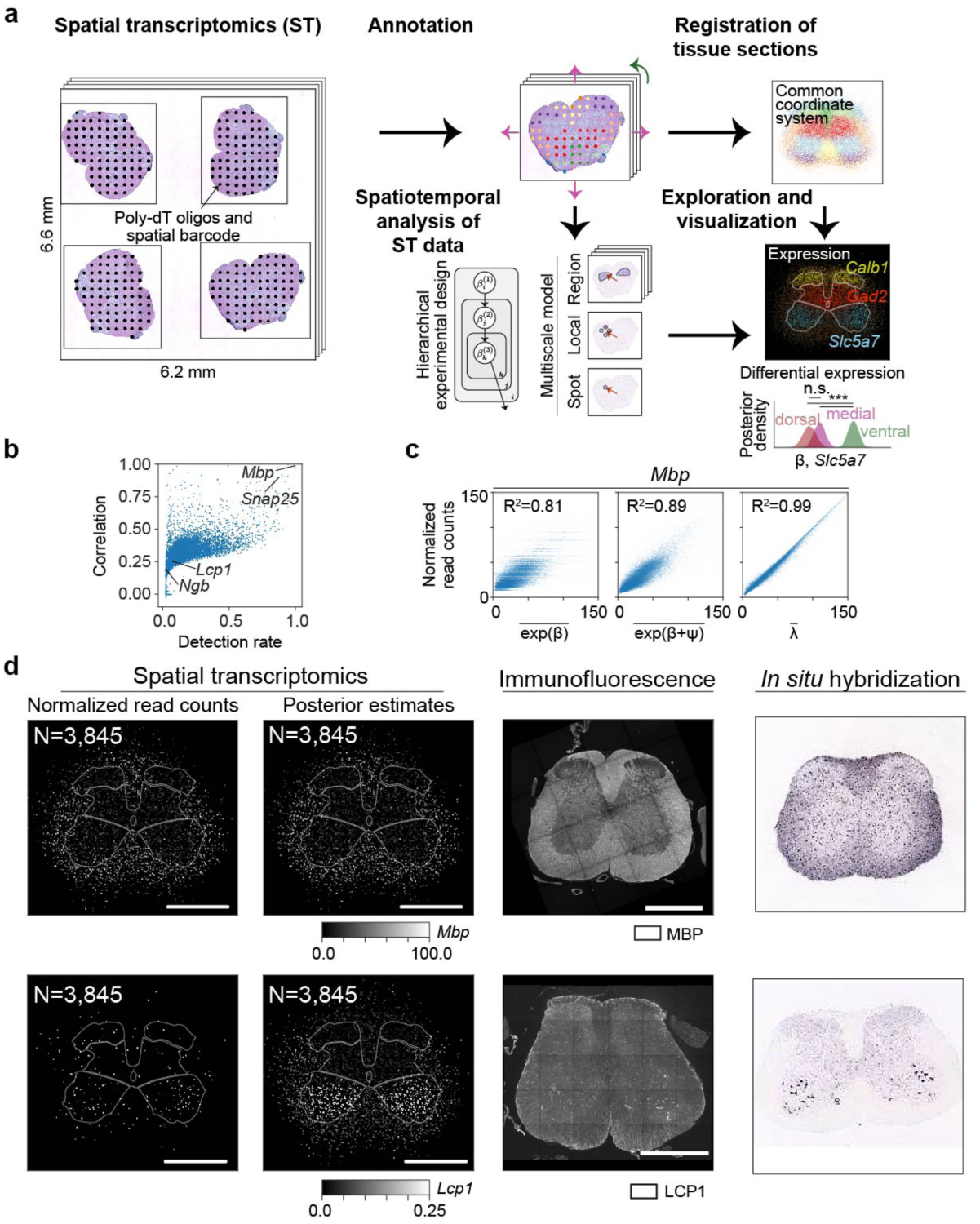
Spatiotemporal modeling of spatial transcriptomics data. (**a**) Illustration of the proposed spatial transcriptomics analysis workflow composed of histological annotation, tissue section registration (top right panel shows registered tissue sections with color coded spot annotations), experimental design aware statistical spatiotemporal data analysis, and visualization steps (lower right panel; spatiotemporal patterns of *Calb1, Gad2*, and *Slc5a7* and quantification of *Slc5a7* expression in anatomical regions [n.s. and *** mark non-significant and significant differences, respectively]). (**b**) The Pearson correlation coefficients of the estimates of the integrative spatiotemporal approach with the estimates obtained using a spatially uninformed approach as a function of detection rate (defined as proportion of non-zero observations). Selected genes have been highlighted. (**c**) Model components’ (region alone 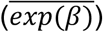 on left, region and local 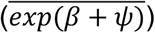 in the middle, and all the three components 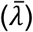 on right; see **Supplementary Fig. 1**) ability to explain variance in the *Mbp* data is studied. The plotted points represent the mouse lumbar spinal cord ST measurements (N=76,136). The coefficients of determination are listed for each case. (**d**) Spatial gene expression of *Mbp* (first row) and *Lcp1* (second row) visualized using the estimates obtained using the spatially uninformed approach (first column) and the integrative spatiotemporal approach (second column) in SOD1-WT lumbar spinal cords at P70 (scale bar is 800 μm). MBP (N = 6 animals) and LCP1 (N = 6 animals) immunofluorescence in SOD1-WT lumbar spinal cords at P70 (third column). *In situ* hybridization images of *Mbp* and *Lcp1* (fourth column) (Image credit: Allen Institute). The number of ST spots is listed.

At its core, our generative model leverages ST data by using hierarchical experimental designs and spatial autocorrelation in measurements at two scales: 1) a tissue context predefined by manually or automatically designated anatomical regions and 2) the local neighborhood encoded automatically by the spatial transcriptomic technology (**Supplementary Fig. 1**; *Methods*). Importantly, the consideration of tissue contexts allows us to share information across tissue sections, enabling detection of reproducible spatiotemporal changes in disease. The choice of distinct tissue contexts, and thereby common coordinate system, is guided by the biological question, spatial resolution, and tissue of interest. To validate Splotch, we analyze a mouse lumbar spinal cord data set^2^ (consisted of 76,136 ST spots across 1,165 tissue sections) and a mouse main olfactory bulb data set^11^ (3,045 ST spots and 12 tissue sections) to demonstrate generalizability. Our spatial model provides a principled way to deal with missing data due to undersampling and, as a result, we are able to quantitate the spatial expression of 11,138 genes on the spinal cord data (and align these genes to the common coordinate), a huge improvement in the much smaller number of transcripts quantified in a single measurement in the median spinal cord ST experiment (1,415 genes and 2,227 unique transcripts per ST spot measurement prior to our Bayesian integration). Moreover, we demonstrate advantages of our approach, especially in the low signal-to-noise ratio (SNR) regime to power downstream analyses described below.

To assess the effect of integrative spatiotemporal modeling, we compared the Splotch estimates with the estimates obtained using a spatially uninformed approach in the spinal cord data (**Fig. 1b**). As expected, in the high SNR regime, both approaches yield similar results as exemplified by myelin basic protein (*Mbp*) and synaptosomal associated protein 25 *(Snap25*) encoding a highly abundant myelinating protein^12^ and a component of the trans-SNAR complex^13^, respectively (**Fig. 1d, Supplementary Fig. 2a**). Spatial (region and local) autocorrelation components of the model capture significant amounts of the variance in the data (**Fig. 1c**), confirming the existence of spatial autocorrelation in ST data, and thus validating our multiscale approach. Importantly, the spatially uninformed approach fails to capture expression localization predicted by Splotch of lowly expressed genes and subsequently confirmed at transcript and protein level, such as lymphocyte cytosolic protein (*Lcp1*) in ventral horn and neuroglobin (*Ngb*) in dorsal horn (**Fig. 1d, Supplementary Fig. 2b-d**). Next, we tested how previous methods^9,10^ for analyzing ST data at the level of individual tissue sections perform on the task of identifying spatially variable genes. Notably, the spatial variability detection by SpatialDE and trendsceek of the four validated genes (*Mbp, Snap25, Lcp1*, and *Ngb*) varies greatly across the tissue sections (**Supplementary Fig. 2a-b**). Although SpatialDE classifies similar spatial variabilility of the genes, that were identified by Splotch, on a subset of the tissue sections (**Supplementary Fig. 2c-d**) it exhibits a dramatically lower detection rate (271 of the 6100 (4.4%) q-values are ≤ 0.05). The low detection rate of SpatialDE and trendsceek is presumably is due to the low SNR and insufficient spatial sampling of physically small (individual mouse) tissue sections. Splotch’s performance is due to the hierarchical and multi-slice integration afforded by the Splotch generative model. Furthermore, we used posterior predictive checking^14^ to validate the model assumptions of Splotch (**Supplementary Fig. 4a**; *Methods*). Reassuringly, the high similarity of the data generated from the posterior predictive distribution with experimental data for selected genes from the high and low SNR regimes demonstrates Splotch’s ability to accurately capture the main characteristics of data-generation process, such as variation and localization (**Supplementary Fig. 4b-f**). Increasing the number of mice improves the estimation independent of the expression level, whereas, increasing the number of tissue sections per mouse improves the estimation of lowly expressed genes (**Supplementary Fig. 5**; *Methods*). Collectively, these results emphasize the merits and robustness of integrative probabilistic modeling of ST data, resulting in increased effective resolution of inferences especially among lowly expressed genes, and thus enabling downstream analyses described below.

To gain insights into transcriptome dynamics of mouse lumbar spinal cord, we quantitate similarities of estimated spatiotemporal expression maps across genes (**Fig. 2a**; *Methods*). Furthermore, we can identify distinct gene sets (modules) that are spatiotemporally co-expressed, visualize modules’ composite spatiotemporal expression maps, and analyze modules’ cell type compositions using independent cell-type level data (*Methods*). To demonstrate the feasibility of this approach, we consider one of the 31 identified modules from our mouse lumbar spinal cord data^2^ (**Supplementary Fig. 6a**). The module is composed of 364 genes showing enrichment of ubiquitin mediated proteolysis and dopaminergic synapse pathways among others^2^. Moreover, these genes are selectively expressed in grey matter with higher expression in medial grey and ventral horn compared to dorsal horn, and show reduced expression in SOD1-G93A lumbar spinal cord at later time points. This module is mainly composed by genes expressed in neurons as 136 genes out of the 364 genes, such as ELAV like RNA binding protein 1 (*Elavl2*) studied in the context of axon degeneration in ALS^15^, are highly expressed in neurons in the light of two independent cell type level data sets^16,17^ (**Supplementary Fig. 6b-d**). The spatiotemporal pattern of *Elavl2* was confirmed at the protein level (**Supplementary Fig. 6e**), indicating that reduced *Elavl2* at later time points reflects the well characterized progressive loss of large *α*-motor neurons of the ventral horn in this mouse model. Taken together, we have described a roadmap for interrogating coordinated transcriptional programs, spatially distinct cell type subpopulations, and interplay between cell types in complex tissues and diseases from the Splotch-based spatiotemporal expression maps.

**Figure 2.**
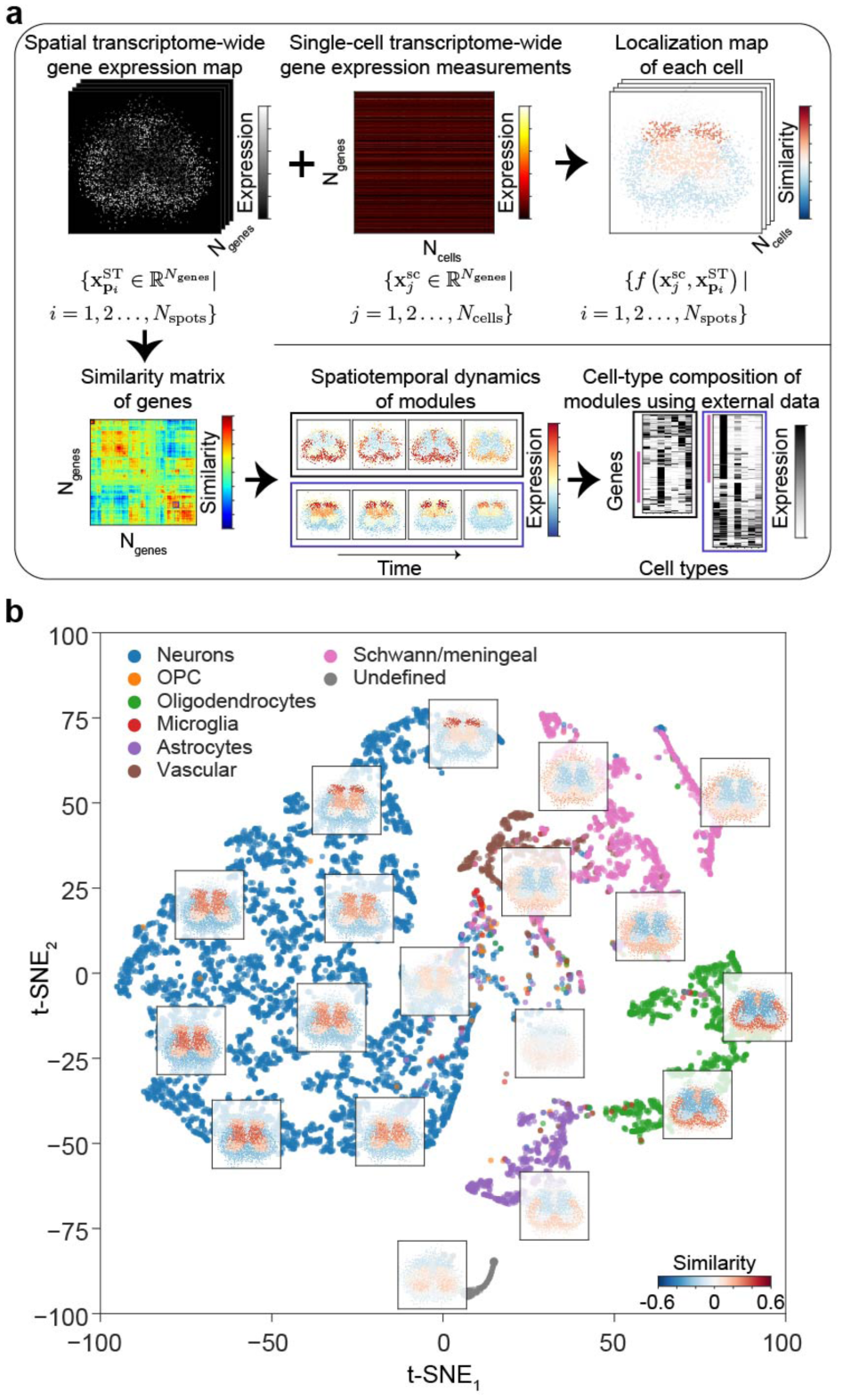
Applications of transcriptome-wide spatial gene expression maps. (**a**) Workflows for unbiased localization of scRNA-seq data using ST data (top) and detecting spatiotemporal transcriptional programs and dissecting corresponding cell types (bottom). (**b**) A visualization of the localization maps of the mouse lumbar spinal cord scRNA-seq data (32,336 cells) using t-SNE. Localization maps are visualized for selected cells. Each localization map is calculated based on 3,845 ST spots. Different colors are used to denote the previously assigned cell type labels^17^.

A limitation of the current single-cell RNA sequencing (scRNA-seq) approaches is the loss of spatial information on tissue dissociation/suspension. Previous attempts to reconstruct spatial localization of scRNA-seq data have used landmark genes with binarized expression to define a basis^18–20^. We hypothesized that the Splotch-based transcriptome-wide spatial expression maps enable the inference of localization of scRNA-seq data without relying on landmark genes (**Fig. 2a**). To test this idea, we considered a recent mouse lumbar spinal cord single nucleus RNA sequencing (snRNA-seq) data set^17^. First, we calculated localization map for each profiled cell using the SOD1-WT (P70) transcriptome-wide spatial map (*Methods*). Dimension reduction of the calculated cell-specific localization maps revealed a continuum of localization maps (**Fig. 2b**); strikingly, the localization map manifold grouped the independently assigned cell type labels^17^ and captured their expected locations in the lumbar spinal cord. The two distinct groups of Schwann/meningeal labeled cells (**Fig. 2b, Supplementary Fig. 7a**) selectively express Schwann and meningeal marker genes^17^ (**Supplementary Fig. 7c**), suggesting that we are able to identify cell localization with high sensitivity. We tested whether we are able to localize regional neuronal subtypes in **Supplementary Fig. 7c**. Our localization results for neuronal cells capture independently assigned^17^ location labels along the dorsal-ventral axis and reflect expression gradient of neuron-expressed genes (**Supplementary Fig. 7c**,**d**). Collectively, these results suggest the feasibility of using ST data to enable unbiased localization of sn/scRNA-seq data and demonstrate the value of using publicly available data in order to combine the complementary strengths of the data types.

Here, we have described computational methods matched to ST technology for interrogating spatiotemporal dynamics of diseases, cell-cell communication, and regulatory dynamics in complex tissues. We have demonstrated the use of Splotch in the context of central nervous system and its dysfunction in amyotrophic lateral sclerosis. Moreover, we analyzed a mouse main olfactory bulb data^11,21^ to test detection of spatial gene expression differences, interrogation of spatial transcriptional programs, and localization of scRNA-seq data (**Supplementary Figs. 8-9**). One potential drawback of our method’s core model is that it could produce overly smooth spatial gene expression maps for lowly expressed genes due to its assumption of spatial autocorrelation. However, our experimental validations show that we are able to detect biologically relevant spatial expression changes even for lowly expressed genes as a result of the integrative and spatiotemporal approach relying on tissue contexts. Increased spatial resolution transcriptome wide measurements would allow us to model smaller tissue contexts while preserving sufficient statistical power. Identification of coordinated spatiotemporal gene expression patterns provides a unique to view the interplay of cell types involved in disease in intact tissue. Localization of scRNA-seq using ST data is particularly useful when studying less characterized tissues lacking information on marker genes. Undoubtedly, we lose some localization accuracy due to the multi-cell nature of ST, and our localization results reflect consensus (as the localization maps are estimated from a large number of tissue section across mice in the datasets analyzed here). In conclusion, we believe the work described here is a valuable component in future attempts to study the functioning and development of complex tissues using multiple possible experimental designs that include spatial transcriptomics.

## Methods

### Analysis of ST data

We index genes (*i*), tissue sections (*j*), and spots (*k*) as follows *i* = 1,2, …, *N*_genes_, *j* = 1,2, …, *N*_tissues_ and 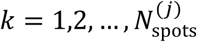. Let *y*_*i, j, k*_ denote the number of UMIs for *i*th gene at *k*th spot on *j*th tissue section. Splotch’s statistical model is a zero-inflated Poisson (ZIP) regression model utilizing a generalized linear model (GLM) with three components to model the rate parameters *λ*_*i,j,k*_: 1) tissue context level component (random variables *β*_*i*,_), 2) local neighborhood level component (random variables *ψ*_*i,j*_), and 3) spot-level component (random variables *ε*_*i,j,k*_) (**Supplementary Fig. 1b**). The hierarchical modeling of the rate parameters of the ZIP regression model enables us to model overdispersion in the observed counts. We use the exponential link function, i.e. 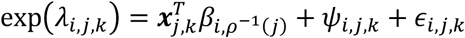 where ***x***_*j,k*_contains one-hot encoded spot annotation information and the function ρ^-1^(*j*) maps the tissue index *j* to its origin in terms of genotype, timepoint, sex, and mouse (**Supplementary Fig. 1a**). Our core statistical model for analyzing ST data was used previously in the context of ST interrogation of ALS^2^. Next, we will describe the model in detail and illustrate model set up in the context of our mouse spinal column data.

The linear model (random variables *β*_*i*,._∈ ℝ^11^) is formulated on 11 tissue contexts and it encodes the hierarchical experimental design (**Supplementary Fig. 1c**). ST spots were annotated based on their location on the tissue using 11 tissue contexts: ventral medial white, ventral lateral white, medial lateral white, dorsal medial white, ventral horn, medial grey, dorsal horn, central canal, ventral edge, lateral edge, and dorsal edge. We explicitly model three different levels of the hierarchical experimental design: 1) *β*_*i*_,_*g*_,_*t*_ (genotype *g* and timepoint *t*), 2) *β*_*i*_,_*g*_,_*t*_,_*s*_ (genotype *g*, timepoint *t*, and sex *s*), and 3) *β*_*i*_,_*g*_,_*t,s,m*_ (genotype *g*, timepoint *t*, sex *s*, and mouse *m*) (**Supplementary Fig. 1a**). Mouse-level parameters *β*_*i*_,_*g*_,_*t,s,m*_ are used to model individual tissue sections from mice. We infer standard deviations at sex 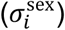 and mouse 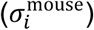 levels. We assume that some of the variation in data is explained by the tissue context of the ST spots; in other words, the linear model captures spatial autocorrelation in terms of tissue contexts. As a result, the linear model and the use of tissue contexts allows us to share information across tissue sections and mice. With this framework for integrating multiple slices/experiments in place, we can study the posterior distributions of the latent parameters *β* at different levels of the hierarchical experimental design, and quantitate expression changes across time, conditions, and tissue contexts. To analyze the mouse main olfactory bulb data set, we annotated the ST spots based on their location on the tissue using five tissue contexts: olfactory nerve layer, glomerular layer, outer plexiform layer, mitral cell layer, and granular cell layer; additionally, we consider two hierarchical levels: genotype and mouse.

Additional spatial autocorrelation at the level of individual tissue section (random variables 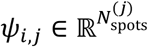) is modeled using the conditional autoregressive (CAR) prior which defines a Markov random field over the ST spots on each ST array

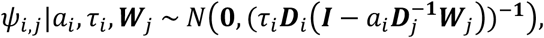

where *a*_*i*_ is a spatial autocorrelation parameter, *τ*_*i*_ is a conditional precision parameter, ***D***_*i*_ is a diagonal matrix (containing the numbers of neighboring ST spots for each ST spot), and ***W***_*j*_ is the adjacency matrix (zero diagonal),

We assume that each ST spot is dependent of ST spots in its local neighborhood defined by the 4-“connectivity” (**Supplementary Fig. 1b**). We infer the level of spatial autocorrelation (*a*_*i*_) and conditional precision (*τ*_*i*_) parameters (**Supplementary Fig. 1c**). Collectively, the random variables 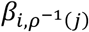 and *ψ*_*i,j*_ capture spatial autocorrelation on two different scales (tissue context and local neighborhood scale respectively).

The third model component (random variables *ε*_*i,j,k*_) captures variation at individual ST spots. We assume that spot-level variations are independent and identically distributed (**Supplementary Fig. 1c**), and we infer their standard deviations (*σ*_*i*_).

We consider the common formulation of the hierarchical zero-inflated Poisson model for counts *y*_*i,j*_,_*k*_

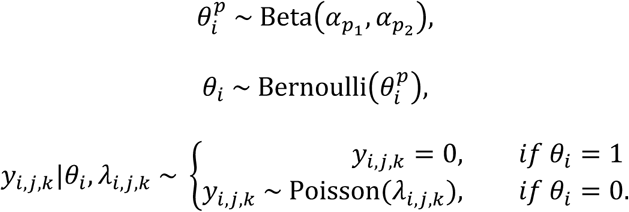

Then, we marginalize the binary variable *θ*_*j*_ (mixture component indicator of Poisson and “zero”), and obtain the ZIP likelihood 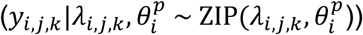

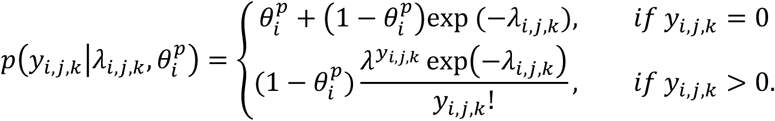

We infer the parameters 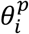.

To adjust for varying sequencing depths across ST spots, we use non-random exposure variables (size factors). That is, for each spot *k* on tissue section *j*, we calculate size factor as *s*_*j,k*_= *N*_*j,k*_ /2208, where *N*_*j,k*_ is the number of UMI counts from the spot *k* on the tissue section *j* (the value 2,208 represents the median sequencing depth for mouse spinal dataset, the same value was used for olfactory bulb to enable direct comparison of *λ* values). Then, for each spot, we adjust/find its rate parameters *λ*_*i,j,k*_ with the size factor *s*_*j,k*_ as *λ*_*i,j,k*_ *s*_*j,k*_ to consider differential sequencing depth. Therefore, our counts *y*_*i,j,k*_ are distributed as follows 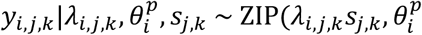

The prior definitions of 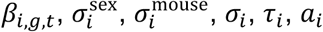, and 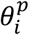 are given in **Supplementary Fig. 1c**. We implemented the model in Stan^22^ and did full Bayesian inference using the adaptive HMC sampler with the default parameters. We sampled four independent chains (each with 500 warm-up and 500 sampling iterations) and monitored the convergence using the R-hat statistic.

It is desirable to have spatially resolved gene expression measurements registered in a common coordinate system to allow comparisons and visualization of spatial gene expression across samples. If the tissue of interest has highly stereotypical architecture, then a common coordinate system can be obtained through registration of histological images using, e.g., computer vision. Whereas, in the cases where the spatial tissue composition varies across samples (e.g. tumor samples) it is less clear how the common coordinate system should be chosen. Here the registration of the spinal cord tissue sections was done by using ventral horn and dorsal horn spots on each tissue section as described previously^2^. Briefly, we first estimated optimal rotations by simultaneously aligning left and right ventral and dorsal horn spots. Second, rotated tissue sections were translated using the centers of mass of ventral and dorsal horn spots. We found that this approach was more robust than the direct registration of the images of the hematoxylin and eosin stained tissues as they showed more variability due to tissue handling and processing. To register the mouse main olfactory bulb tissue sections, we first estimated optimal rotations by aligning left and right granular cell layer spots, then the rotated tissue sections were translated using the centers of mass of granual cell layer spots.

Detection of differential expression between conditions and regions was done by quantitating element-wise differences in the estimated posterior distributions of the random variables *β*_*i*_ using the Savage-Dickey density ratio as described previously^2^.

### Spatially-unaware analysis of ST data

First, the UMI count of each gene per ST spot *y*_*i,j,k*_ is divided by the total number of UMIs at that spot *N*_*j,k*_, and then multiplied by the median total number of UMIs across ST spots (2,208 for mouse spinal dataset). This procedure corresponds to the exposure procedure through size factors as described above and makes the estimates of the two approaches comparable.

### Comparison of Splotch with SpatialDE and trendsceek

SpatialDE^9^ and trendsceek^10^ have been previously proposed for identifying spatial gene expression trends from data in an unsupervised manner. Splotch differs from SpatialDE and trendsceek in the following ways: 1) Splotch provides a way to quantify expression differences between conditions and anatomical regions etc., 2) its statistical model is tailored for count data, and thus deals with uncertainty of low counts rigorously, and 3) it analyzes multiple tissue sections simultaneously and takes into the experimental design in order to quantify biological and experimental variation at different levels.

The latest version of SpatialDE (v1.1.0) was used. First, the unfiltered counts are transformed (norm_expr = NaiveDE.stabilize(counts.T).T) and normalized (resid_expr = NaiveDE.regress_out(sample_info, norm_expr.T, ‘np.log(total_counts)’).T). Then, SpatialDE is used to analyze the normalized data (results = SpatialDE.run(X, resid_expr), where X contains the original spot coordinates) resulting in q-values and p-values used in the comparison. The same procedure is done for every tissue section separately and results are pooled.

The latest version trendsceek (v.1.0.0) was used. First, we discarded lowly expressed genes (counts = genefilter_exprmat(counts, 1, 3)) because trendsceek fails to analyze them. Second, we added small uniform jitter (U(−0.3,0.3)) to the spot coordinates to ensure that the summary statistics calculation will not fail. Third, the counts are normalized (counts_norm = deseq_norm(counts, 1)) and the marks are set (set_marks(pos2pp(coordinates), counts_norm, log.fcn = log10)). Finally, trendsceek is used to the analyze the data (trendstat_list = trendsceek_test(pp, 1000, 1)) and the obtained p-values are used in the comparisons presented. The same procedure is done for every tissue section separately and results are pooled.

### Spatiotemporal and disease-dependent co-expression analysis

After running the integrative model described above (and illustrated in **Fig. 1a**), we collect all resulting posterior means of the spot-level rate parameters *λ*_*i,j,k*_ in a 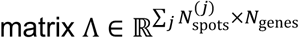 (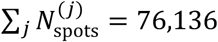 and *N*_genes_ = 11,138, for mouse spinal cord data set; 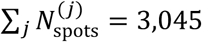 and *N*_genes_ = 13,340, for mouse main olfactory bulb data set). Then, we calculate the matrix 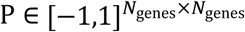 that contains Pearson correlation coefficients for each pair of genes using the matrix Λ. To find groups of genes that have similar spatiotemporal dynamics, we used hierarchical cluster analysis (L1 norm and average linkage) on P and set the distance threshold so that the main diagonal blocks would be separated, resulting in 31 and 27 co-expression modules on mouse spinal cord^2^ and olfactory bulb data sets, respectively.

To calculate consensus spatiotemporal expression map of the *o*^th^ co-expression module, we first take a submatrix of Λ containing the columns that correspond to the genes of the *o*th coexpression module, second, we standardize the columns of the submatrix to transform genes’ expression values to similar scale, and third, we calculate the averages over the rows to summarize co-expression module’s expression at each spot. Given that each spot is associated with genotype, timepoint, and coordinates in the common coordinate system, we can visualize the calculated consensus spatiotemporal map of each co-expression module^2^.

The initial analysis of the identified co-expression modules on mouse spinal cord data suggested that each of them was composed by multiple cell types, leading to a need of deconvoluting the cell types contributions. To identify the cell-type components in the co-expression modules, we utilized published cell-type level expression data as described previously^2^. Briefly, we detected distinct expression patterns of the genes of each co-expression module based on cell-type level bulk or single-cell RNA sequencing data using hierarchical clustering, leading to submodules. Then, we detected which of the identified submodules are showing cell-type specific expression, i.e., are the genes of the given submodule specifically expressed in astrocytes.

### In situ hybridization images

The *in situ* hybridization images of *Mbp, Lcp1, Snap25*, and *Ngb* included in Fig. 1 and Supplementary Fig. 2 were downloaded from http://mousespinal.brain-map.org (Image credit: Allen Institute):

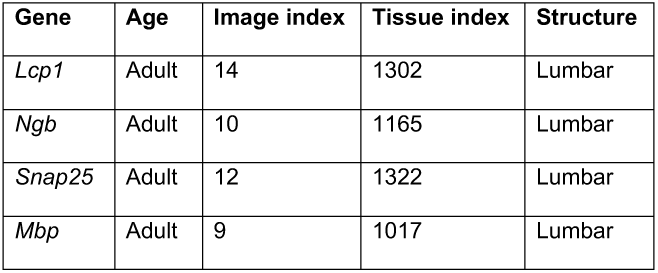

| Gene | Age | Image index | Tissue index | Structure |
| --- | --- | --- | --- | --- |
| *Lcp1* | Adult | 14 | 1302 | Lumbar |
| *Ngb* | Adult | 10 | 1165 | Lumbar |
| *Snap25* | Adult | 12 | 1322 | Lumbar |
| *Mbp* | Adult | 9 | 1017 | Lumbar |

### Posterior predictive checking

We evaluate our overall model fit to data using posterior predictive checking as follows; we first describe the method we use for PPC and then describe its adaptation in the context of Splotch. Let us consider a general case to define posterior predictive distribution: let ***Y*** denote the observed data, *y*^new^ denote the unobserved data, *θ* denote the parameters, *α* denote the hyperparameters. The posterior predictive distribution is the distribution of unobserved values *y*^new^ conditional on the observed values ***Y***

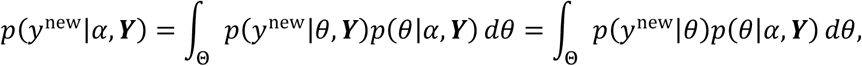

where the last equality follows from the assumption that the observed and und unobserved data are conditionally independent given *θ*. Whereas, the prior predictive distribution is

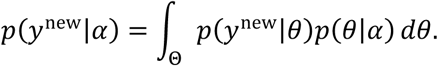

Next, we will describe how we implemented posterior predictive checking in the context of Splotch and ST. First, we consider an *in silico* ST array with similar layout as actual ST slides (**Supplementary Fig. 4a**). Second, we overlay all the registered ST spots on the *in silico* ST array and discard those *in silico* ST array spots that have low coverage by the registered ST spots (less than 50 spots within 0.2 units in terms of Euclidean distance) (**Supplementary Fig. 4a**). Third, we automatically annotate each of the *in silico* ST array spots by taking the most abundant annotation tag in the spot neighborhood (0.2 units in terms of Euclidean distance) (**Supplementary Fig. 4a**). Fourth, we sample data from the posterior predictive distribution in the generated quantities block using the posterior samples of 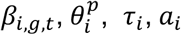, and *σ*_*i*_. While sampling from the posterior predictive distribution we assume that the sequencing depth of each spot on the *in silico* array is 2,208, and thus the corresponding size factors are 1 (**Supplementary Fig. 4b-e**). In more detail, each iteration, we first calculate 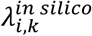 using the iteration-specific samples of *β*_*i,g,t*,_ *τ*_*i*_ (used to draw 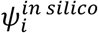), *a*_*i*_ (used to draw 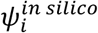), and *σ*_*i*_ (used to draw 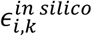) and the corresponding spot annotation 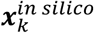, and second, we generate a single count for each spot from the ZIP distribution 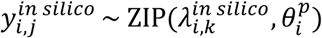, where 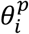 is the current sample.

Samples from the prior predictive distributions (**Supplementary Fig. 4f**) were generated similarly with one important exception: the used parameters values were sampled from the prior instead of from the posterior.

### Subsampling of data and its effect on estimation

We chose to use the ST data of SOD1-G93A mice at P70 as the data set was composed of altogether 95 tissue sections (8, 5, 7, 21, 14, 12, 4, and 24 tissue section per mouse) collected from eight mice (four males and four females). To subsample the data, we randomly selected the given number of mice and then for each of the selected mice we randomly selected the given number of tissue section per mouse (if the selected mouse had less than the given number of tissue sections, then we took all the tissue sections of that mouse). Then, we analyzed each of the subsampled data set (the twelve combinations of 1, 2, 4, and 8 mice with 2, 4, and 8 tissue sections per mouse) and the full data set (95 tissue sections and eight mouse) separately.

To compare the estimated posterior distributions of *β*_*i*,SOD1-G93A,P70_ at the gene and tissue context levels, we first calculated the posterior means and their standard deviations, which were used to obtain normal distribution approximations of the posterior distributions. Third, we used the Kullback-Leibler divergence to quantitate the differences between the normal distributions. In more detail, we quantitate how much information we lose by using subsampled data sets (with respect to the full data set).

We studied the effect of number of mice and number of tissues section per mouse on the spatial gene expression estimation (**Supplementary Fig. 5**). Even with single mouse and two tissue sections, we already capture meaningful signal, but the estimation accuracy improves significantly with four mice (**Supplementary Fig. 5a**). Increase in the number of mice improves the estimation independent of the expression level, whereas, increasing the number of tissue sections per mouse improves the estimation of lowly expressed genes (from four to eight per mouse) (**Supplementary Fig. 5b**). These numbers clearly depend on the sequencing depth and the biological system of interest, for instance, in the case of human samples the number of individuals has to be increased due to the intrinsic biological variation.

### Localizing scRNA-seq data

We obtained the count table 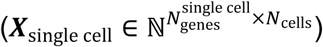 representing the snRNA-seq based data of adult mouse lumbar spinal cord generated by Sathymurthy *et alii*^17^. The computationally derived cell-type labels associated with the cells were kindly provided by the authors. First, we impute sRNA-seq data matrix ***X***_single cell_ using MAGIC^23^ to obtain 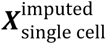 (filter_scseq_data(filter_cell_min=1000, filter_cell_max=100000, filter_gene_nonzero=None, and filter_gene_mols=None) and run_magic(n_pca_components=20, random_pca=True, t=None, compute_t_make_plots=True, t_max=12, compute_t_n_genes=500, k=30, ka=10, epsilon=1, rescale_percent=99)). We consider the SOD1-WT lumbar spinal cord (P70) ST data as it is the closest match to experimental conditions of the snRNA-seq data (8-12 weeks). That is, we construct a matrix using rate parameters *λ*_*i,j,k*_ (corresponding to SOD1-WT at P70 and 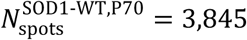

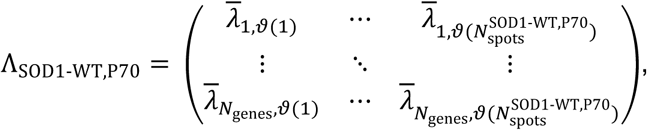

where the function *ϑ*(·) maps running spot indices to tissue and spot indices corresponding to the condition SOD1-WT at P70 and the overbar denotes posterior mean. Then, we standardize the 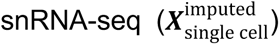 and ST (Λ_SOD1-WT, P70_) data matrices separately across cells (*N*_cells_ = 32,336) and spots within genes, respectively, and consider the common genes (*N*_common genes_ = 10,681), resulting in matrices 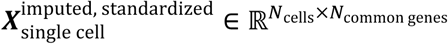 and 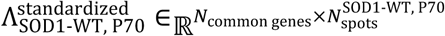. Finally, for each cell we calculate its similarity to each spot by calculating Pearson correlation between its standardized and imputed expression vector (columns of 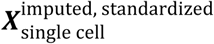) and spots’ expression vectors (columns of 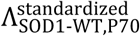), resulting cell-specific similarity vectors that can be visualized in the common coordinate system using the registered spot coordinates (**Fig. 2a**). The same procedure was used with the mouse main olfactory ST and scRNA-seq data sets generated by Ståhl *et alii* and Tepe *et alii*^11,21^, respectively, resulting in 33,107 cells, 12,628 common genes, and 3,045 ST spots.

To visualize the estimated similarity maps, we use t-SNE as implemented in scikit-learn (sklearn.manifold.TSNE) using the PCA initialization (default parameters otherwise). We derive t-SNE maps of the similarity vectors either of all the cells (**Fig. 2b, Supplementary Fig. 7a**) or all the neuronal cells (**Supplementary Fig. 7c**). Similarity maps of randomly selected cells are visualized. Finally, we overlay the computationally derived cell-type labels associated with the cells on the t-SNE plots.

### Murine ALS models

B6SJLSOD1-G93A transgenic and SOD1-WT transgenic mice were obtained from Jackson Laboratories (Bar Harbor, ME), and maintained in full-barrier facilities at Columbia University Medical Center in accordance with ethical guidelines established and monitored by Columbia University Medical Center’s Institutional Animal Care and Use Committee. SOD1-G93A mice were monitored closely for onset of disease symptoms, including hindlimb weakness and weight loss. Disease end-stage was defined as the inability to become upright in 15s after being placed on their back.

### Spinal cord collections and sectioning

Mice were transcardially perfused with 1X Phosphate buffered saline (PBS) followed by 4% paraformaldehyde (PFA) in 1X PBS. Spinal cords were then dissected and the L3-L5 lumbar region isolated based upon ventral root anatomy. The isolated spinal cord was then post fixed in 4% PFA in 1X PBS for 24hrs at 4°, and then cryoprotected by saturation with 30% sucrose diluted in 1X PBS at 4°. Samples were then embedded in Optimal Cutting Temperature (OCT, Fisher Healthcare, USA), plunged into a bath of dry ice and pre-chilled ethanol until freezing, and stored at −80°C. Cryosections were cut at 10μm thickness onto Superfrost plus slides (VWR International, USA). For LCP1, MBP, and ELAVL2, sections were blocked in 1X PBS supplemented with 5% donkey serum (Jackson Immunoresearch, USA), 0.5% Bovine Serum Albumin (BSA, Sigma Aldrich, USA) and 0.2% Triton X-100 (Sigma-Aldrich, USA) for 1h at room temperature. This was followed by primary antibody staining (diluted in the Triton X-100 blocking buffer) at 4°C overnight, washing in 1X PBS with 0.2% Triton X-100 (PBS-T), and then secondary antibody incubation at room temperature for 4h (1:500 dilution in the Triton X-100 blocking buffer) and washed in PBS-T. For SNAP25 and NGB, sections were blocked in 1X PBS supplemented with 5% donkey serum (Jackson Immunoresearch, USA), 0.5% Bovine Serum Albumin (BSA, Sigma Aldrich, USA) and 0.1% saponin (Sigma-Aldrich, USA) for 1h at room temperature. This was followed by primary antibody staining (diluted in the saponin blocking buffer) at 4°C overnight, washing in 1X PBS with 0.1% saponin (PBS-S), and then secondary antibody incubation at room temperature for 4h (1:500 dilution in the saponin blocking buffer) and washed in PBS-S and subsequently in 1X PBS. The slides were mounted in Vectashield (Vector Laboratories, USA) and cover slipped (VWR, USA). Primary antibodies were diluted as follows: MBP (Abcam; Ab209328; 1:1000), LCP1 (Atlas Antibodies; HPA019493; 1:250), ELAVL2 (Atlas Antibodies; HPA063001; 1:100), SNAP25 (Atlas Antibodies; HPA001830; 1:250), NGB (Novus; NBP2-48769;1:250). Secondary antibodies were Alexa Fluor 488 or 633 dye conjugated and obtained from Jackson ImmunoResearch.

### Equipment and settings

Confocal images were acquired on a Zeiss LSM 780 with a 20x/0.8 Plan-APOCHROMAT objective (Carl Zeiss Microscopy, Germany). All images were acquired at a x/y resolution of 0.692 microns/pixel, and at a 14bit depth. 488nm and 633nm laser excitation was used with light path settings appropriate for Alexa Fluor 488 and 633 dye emission spectra.

## Data availability

The accession numbers of the used data sets are: GSE120374, PRJNA316587, GSE103892, GSE121891, and GSE52564.

### Code availability

Splotch is available at: https://github.com/tare/Splotch

## Supplementary Figure legends

**Supplementary Figure 1.**
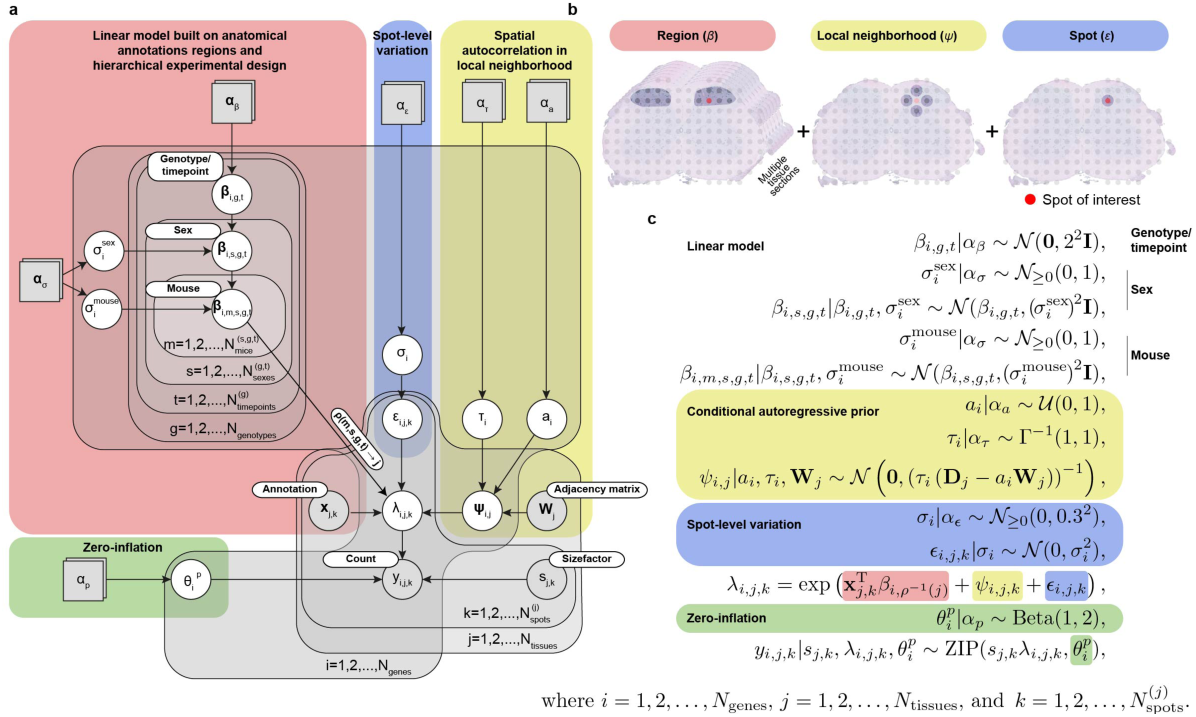
Statistical model. (**a**) Graphical representation of Splotch’s statistical model using plate notation. Grey and white circles represent observable and latent random variables, respectively. Grey squares represent variables with fixed values (user-defined). Parameter names are listed in the circles and squares. Grey plates represent repetitions described in the lower right corners. The red, blue, yellow, and green shaded areas are used to distinguish model components. (**b**) A schematic illustrating the used multiscale model for variance decomposition. (**c**) The distributions of the random variables of the statistical model illustrated in (**a**) are listed. The values α_β_, α_σ_, α_τ_, α_a_, α_ε_, and α_p_, that define the priors of 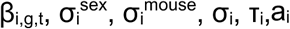, and 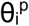 are listed. The red, blue, yellow, and green shaded areas match of (**a**).

**Supplementary Figure 2.**
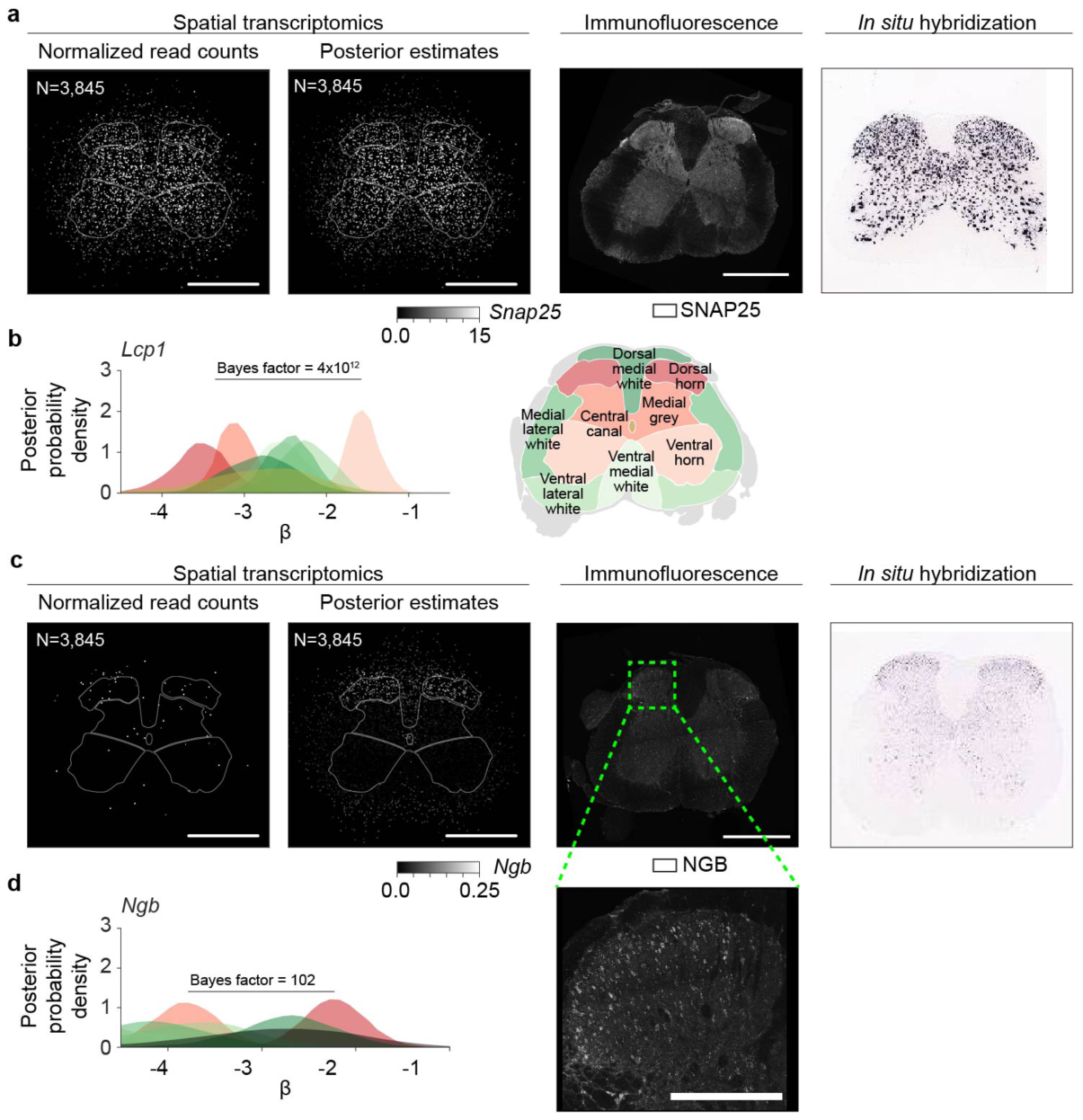
Spatial gene expression in SOD1-WT lumbar spinal cord at P70. (**a**) Spatial gene expression of *Snap25* visualized using the estimates obtained using the spatially uninformed approach (first column) and the integrative spatiotemporal approach (second column) in SOD1-WT lumbar spinal cords at P70 (scale bar is 800 μm). SNAP25 immunofluorescence in SOD1-WT lumbar spinal cords at P70 (third column) (N = 6 animals). *In situ* hybridization image of *Snap25* (fourth column) (Image credit: Allen Institute). The number of ST spots is listed. (**b**) The posterior distributions of the region-specific coefficient parameters β of *Lcp1* in SOD1-WT at P70 (left) (ventral, lateral, and dorsal edge anatomical annotation regions are not considered). The color key is illustrated on right. The Bayes factor to quantify the difference between ventral horn and the combination of medial grey and dorsal horn is listed. (**c**) Spatial expression of *Ngb* visualized as in (**a**) (N = 4 animals). A zoomed view of the dorsal horn (the dashed rectangle) is visualized in lower right (scale bar is 250 μm). (**d**) The posterior distributions of the region-specific coefficient parameters β of *Ngb* in SOD1-WT at P70 visualized as in (**b**). The Bayes factor to quantify the difference between dorsal horn and the combination of medial grey and ventral horn is listed.

**Supplementary Figure 3.**
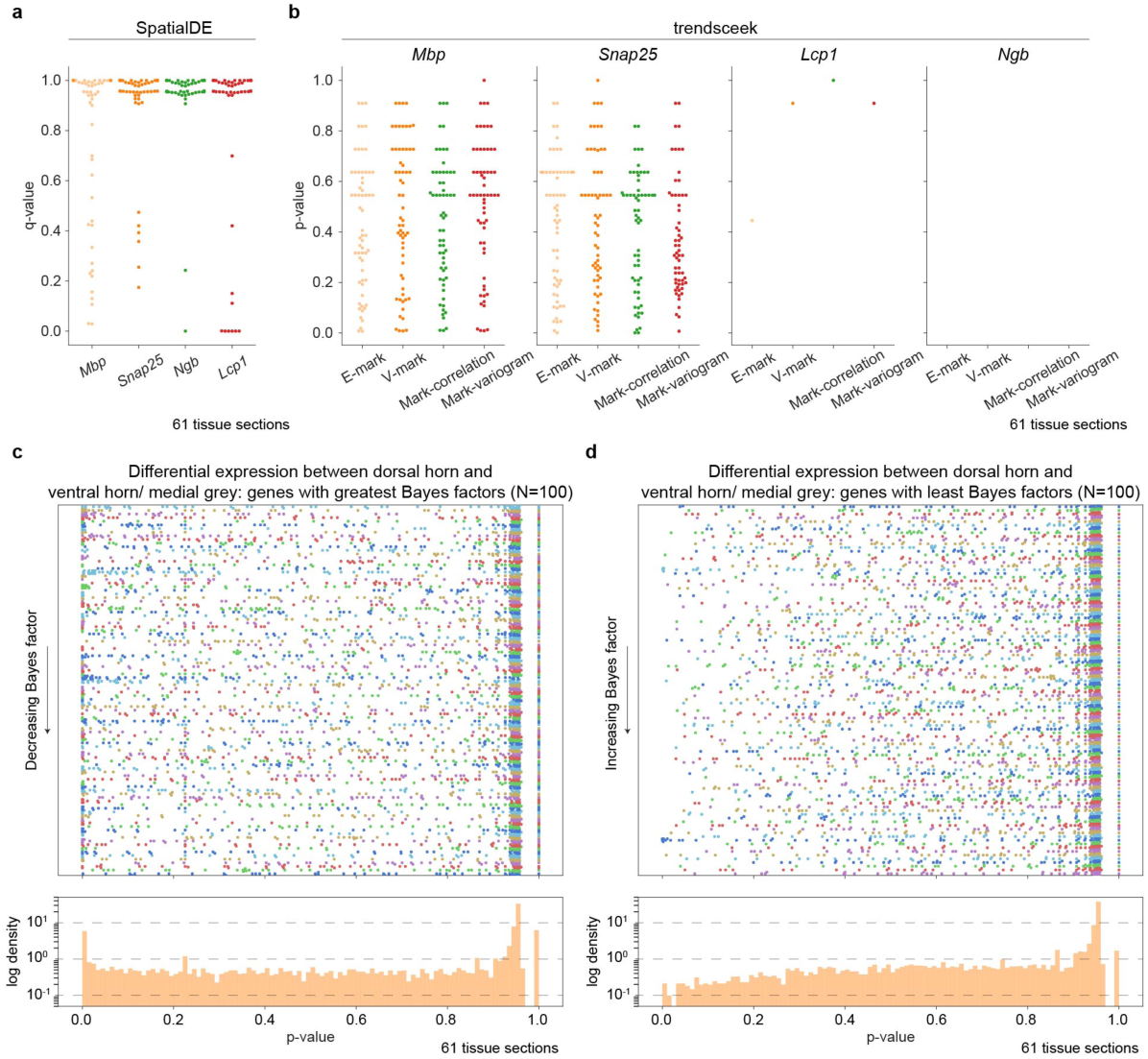
Comparison of Splotch, SpatialDE and trendsceek on SOD1-WT lumbar spinal cord tissue sections at P70. (**a**) Q-values (on y-axis) of *Mbp, Snap25, Lcp1*, and *Ngb* (on x-axis) from the spatial variability test as implemented in SpatialDE are visualized. Each spot represents a tissue section (N=61). Few of these genes show significant differential or spatial differential expression with either tool. These genes were validated by us (with microscopy or via prior work) and are robustly detected with the correct spatial pattern by Splotch. (**b**) Unadjusted p-values (on y axis) from the E-mark, V-mark, mark-correlation, and mark-variogram tests (on x axis) for *Mbp, Snap25, Lcp1*, and *Ngb* (on columns) as implemented in trendsceek are visualized. Each spot represents a tissue section. For each considered gene, those tissue sections without enough sequencing depth for trendsceek are not analyzed (*Methods*). For instance, only one tissue section has enough sequencing depth for *Lcp1*. In all slices/experiments Splotch is able to recover expression estimates for these genes. (**c**) P-value (unadjusted) distributions produced by SpatialDE of all genes identified by Splotch to be differentially expressed in dorsal horn when compared with ventral horn and medial grey (100 genes with the greatest Bayes factors). The genes (rows) have been ordered by Bayes factors (y-axis). The points represent distinct tissue sections. Different colors are used to distinguish adjacent rows (i.e. genes). Additionally, the normalized histogram of p-values across the genes and tissue sections is visualized below. (**d**) As in (**c**) but here we focus on the genes (N=100) with the least Bayes factors; that is, Splotch suggests that these are not differentially expressed in dorsal horn compared to ventral horn and medial grey.

**Supplementary Figure 4.**
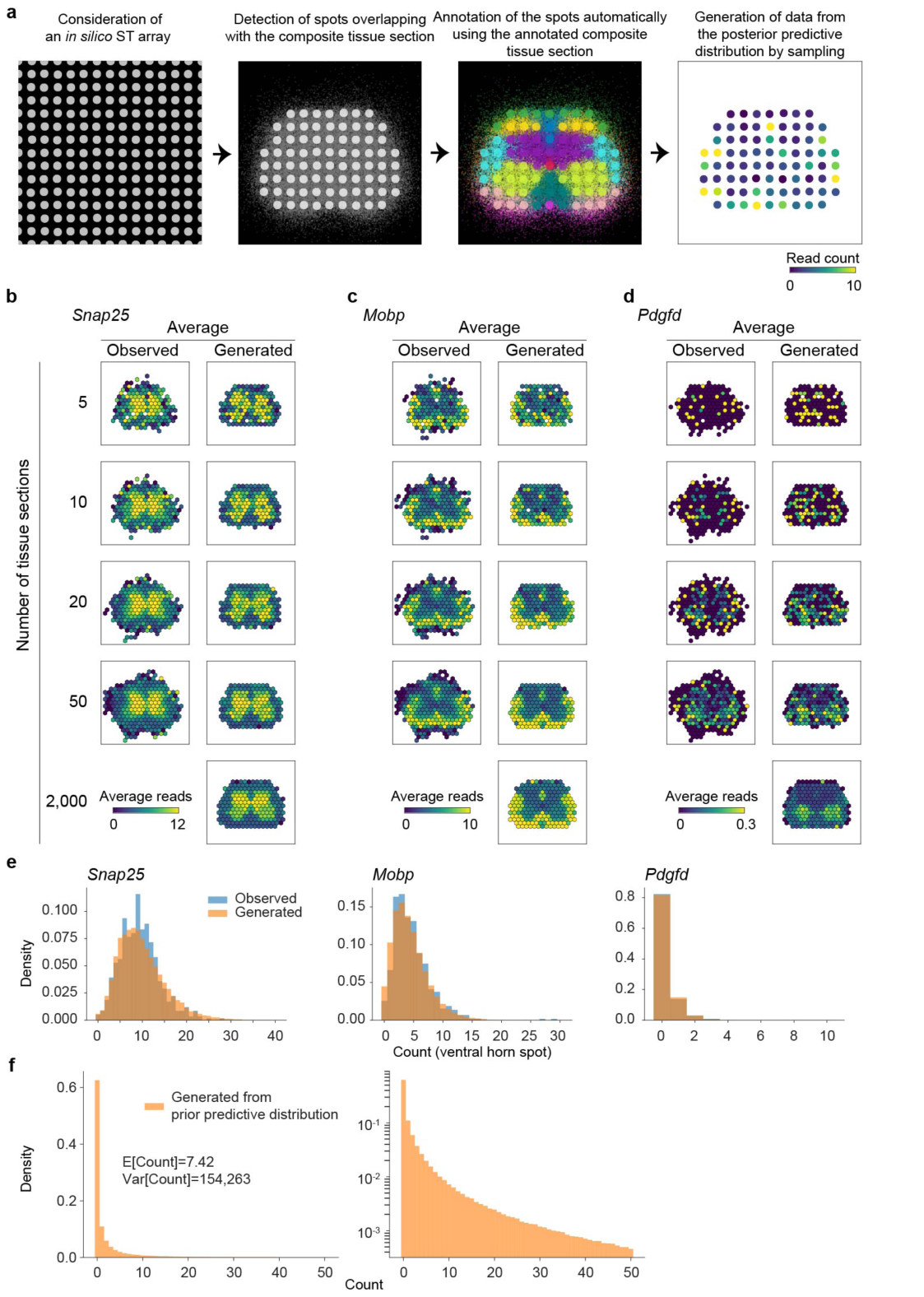
Posterior predictive checks for model validation. (**a**) Workflow of the posterior predictive checking is illustrated. First, an *in silico* ST array is considered and overlaid onto the composite tissue section. Second, those spots not covering the composite tissue section are discarded. Third, the spots are annotated automatically using the annotation information of the composite tissue section (annotation regions are individually color coded). Finally, we generate data from the posterior predictive distribution. (**b**) The posterior predictive checking of *Snap25*. The average expression map calculated based on the observed UMI counts (left column) using either five (first row), 10 (second row), 20 (third row), or 50 (fourth row) randomly selected SOD1-WT lumbar spinal cord tissue sections at P70. The observed counts have been normalized to match the median experiment, i.e. they have divided by the total number of UMIs per spot and multiplied by the median sequencing depth (2,208). The average expression map calculated based on the counts sampled from the posterior predictive distribution assuming the median sequencing depth (2,208) (right column) using either five (first row), 10 (second row), 20 (third row), 50 (fourth row), or 2,000 (fifth row) tissue sections. Small random jitter is added to the coordinates of each simulated tissue section to ensure that we get more uniform coverage. The β parameter used in the simulation corresponds to SOD1-WT lumbar spinal cord tissue sections at P70. (**c**) The posterior predictive checking of *Mobp* as in (**b**). (**d**) The posterior predictive checking of *Pdgfd* as in (**b**). (**e**) Estimated probability mass functions of counts at ventral horn annotated spots in SOD1-G93A lumbar spinal cord at P70 for *Snap25, Mobp*, and *Pdgfd* based on the observed data (blue) and data generated from the posterior predictive distribution (orange) assuming the median sequencing depth (2,208). The observed counts have been normalized to match the median experiment, i.e. they have divided by the total number of UMIs per spot and multiplied by the median sequencing depth (2,208). (**f**) Estimated probability mass functions (linear and log scales on left and right, respectively) of counts based on the data generated from the prior predictive distribution assuming the median sequencing depth (2,208 UMI counts). Sample mean and variance are listed.

**Supplementary Figure 5.**
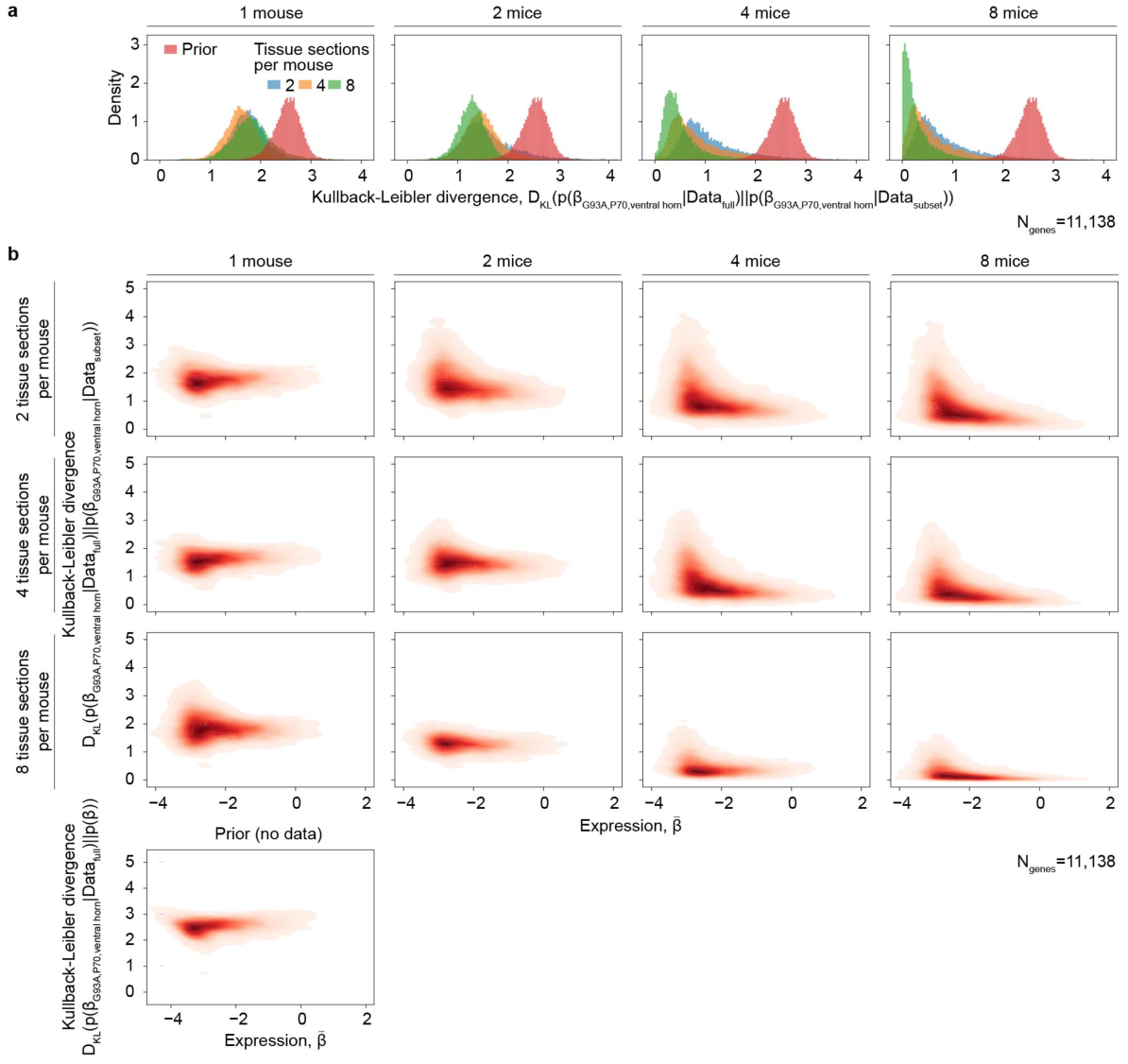
Effect of experimental design on estimation. (**a**) Comparison of the coefficient distributions (SOD1-G93A, P70, ventral horn) across genes (N=11,138) estimated from full (8 mice and 95 tissue sections) and subsampled data. The number of mice and the number of tissue sections per mouse are varied. The difference of the distributions is quantitated using the Kullback-Leibler divergence, that is we quantify the divergence of the distribution estimated from the full data set with respect to the distribution estimated from the subsampled data (how much information is lost when using the distribution estimated from the subsampled data). (**b**) As in (**a**) but here we additionally study the divergences as a function of the coefficient value (posterior mean of the distribution estimated from the full data set).

**Supplementary Figure 6.**
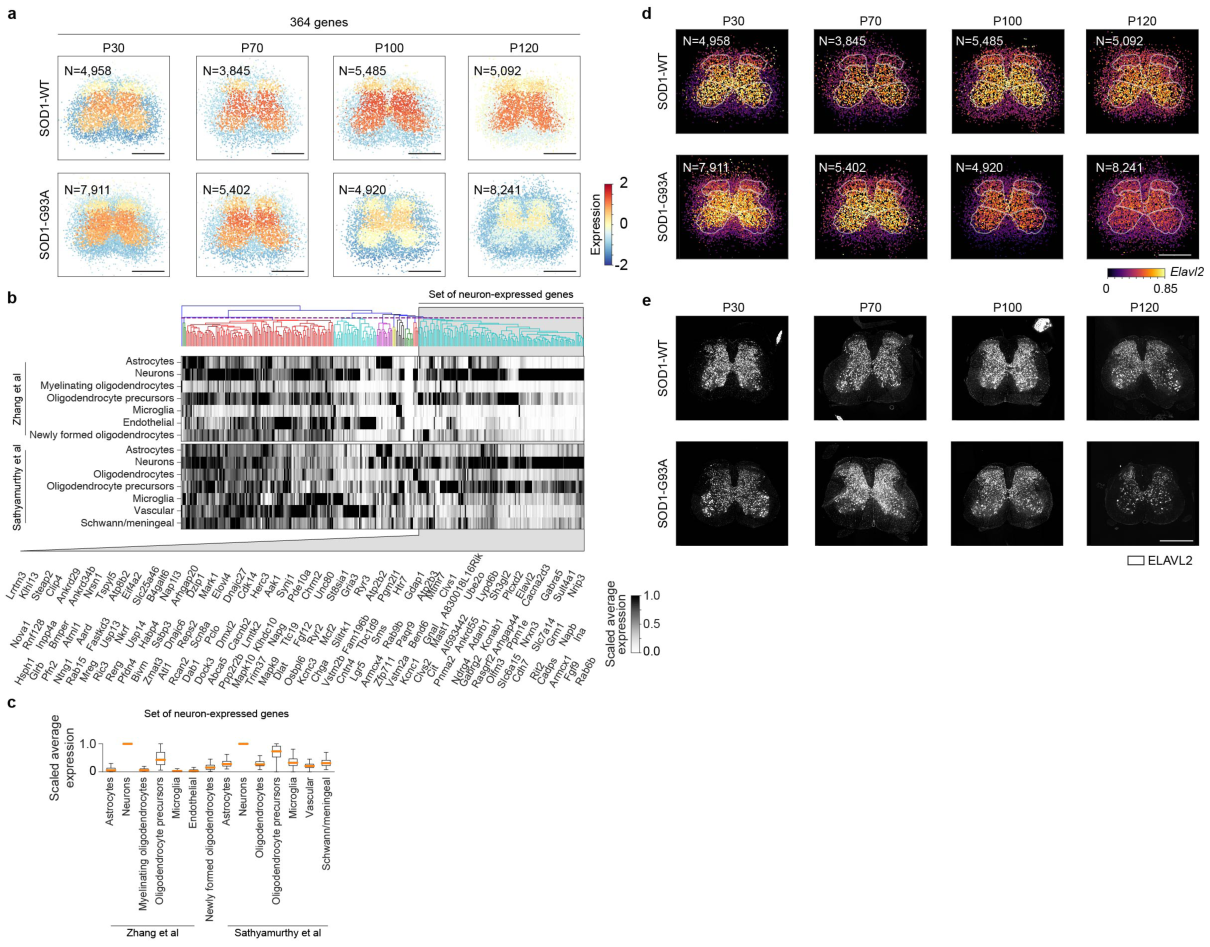
Cell-type composition of co-expression modules. (**a**) Spatiotemporal dynamics of a selected co-expression module containing 364 genes in SOD1-WT and SOD1-G93A lumbar spinal cords at P30, P70, P100, and P120 are visualized. The colors represent averages (across genes) of standardized expressions (across spots within genes) at each spot. The number of ST spots per condition is listed. Scale bar is 800 μm. (**b**) Expression pattern of the genes of co-expression module visualized in (**a**) (top). The genes have been clustered using independent cell-type level gene expression data. The heatmaps show clustered scaled expression data of brain cell types^16^ (top heatmap) and spinal cord cell types^17^ (bottom heatmap). The dashed vertical line in the dendrogram denotes the cutting point. A set of genes that was identified to be enriched of neuron-expressed genes has been highlighted and the gene symbols are listed (bottom). (**c**) The boxplots (rectangle spans first quartile [Q1] to the third quartile [Q3], a line inside the rectangle shows the median [Q2], and the whiskers span from Q1-1.5×(Q3-Q1) to Q3+1.5×(Q3-Q1)) show scaled expression data of the highlighted neuron-expressed genes highlighted in (**b**) in brain and spinal cord cell types. Outliers are not depicted. (**d**) Spatial gene expression of *Elavl2* in SOD1-WT and SOD1-G93A lumbar spinal cords at P30, P70, P100, and P120 are visualized. The number of ST spots per condition is listed. Scale bar is 800 μm. (**e**) ELAVL2 immunofluorescence in SOD1-WT and SOD1-G93A lumbar spinal cords at P30, P70, P100, and P120 (N = 4 animals each). Scale bar is 800 μm.

**Supplementary Figure 7.**
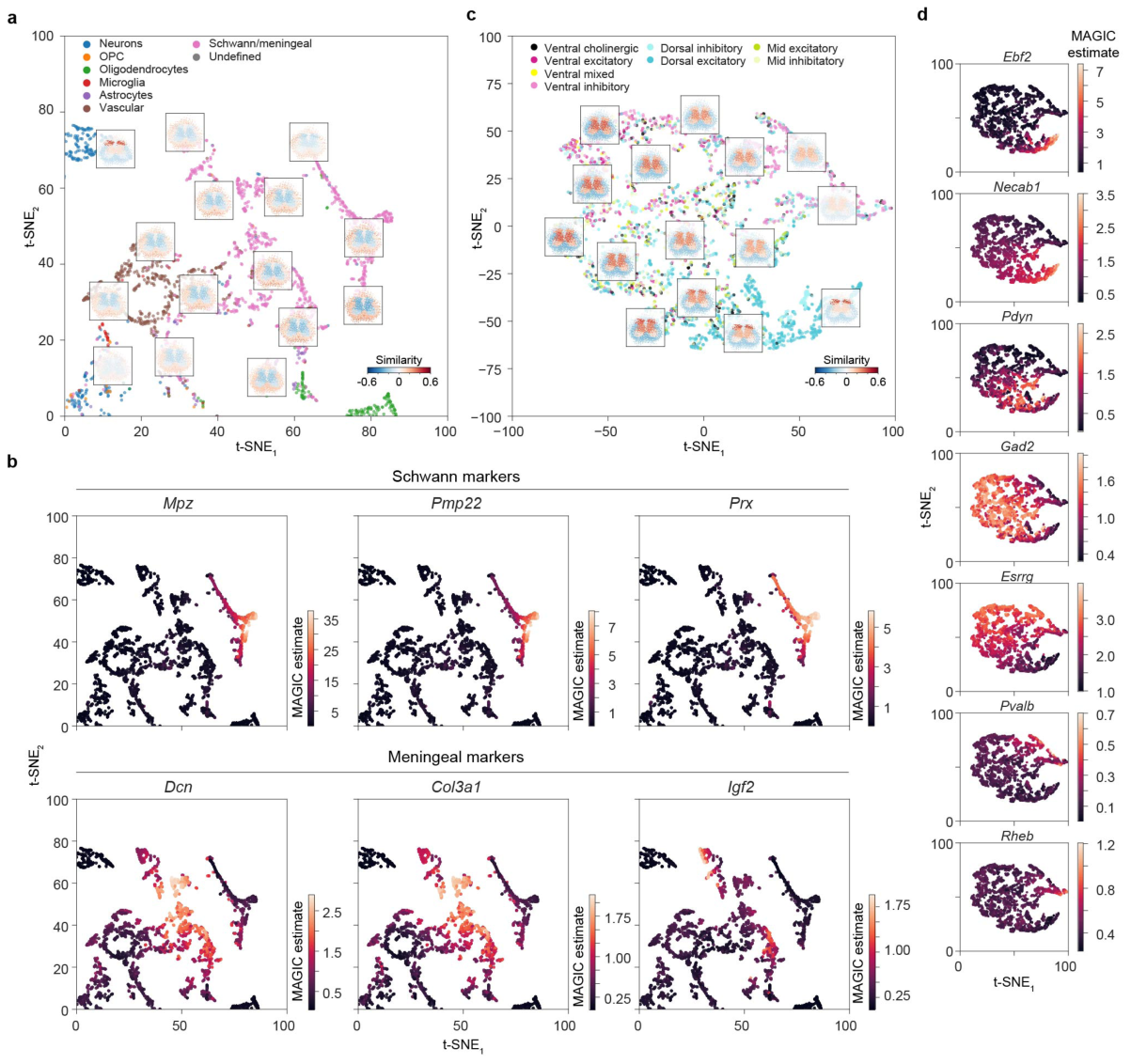
Localization of Schwann and meningeal cells, and neurons. (**a**) The top right quadrant of **Fig. 2b** is visualized. (**b**) Expression of Schwann and meningeal cell marker genes in the Schwann/meningeal labeled cells is visualized. The cells are plotted in the t-SNE space calculated based on the localization maps visualized in (**a**). (**c**) A visualization of the localization maps of the neuronal cells based on the mouse lumbar spinal cord snRNA-seq data using t-SNE (the estimated manifold is different from **Fig. 2b** as only neuronal cells are considered). Localization maps are visualized for selected neuronal cells. Different colors are used to denote the independently assigned location and type labels. (**d**) Expression of *Ebf2, Necab1, Pdyn, Gad2, Pvalb*, and *Rheb* in the neuron labeled cells is visualized. The cells are plotted in the t-SNE space in (**c**) calculated based on the localization maps.

**Supplementary Figure 8.**
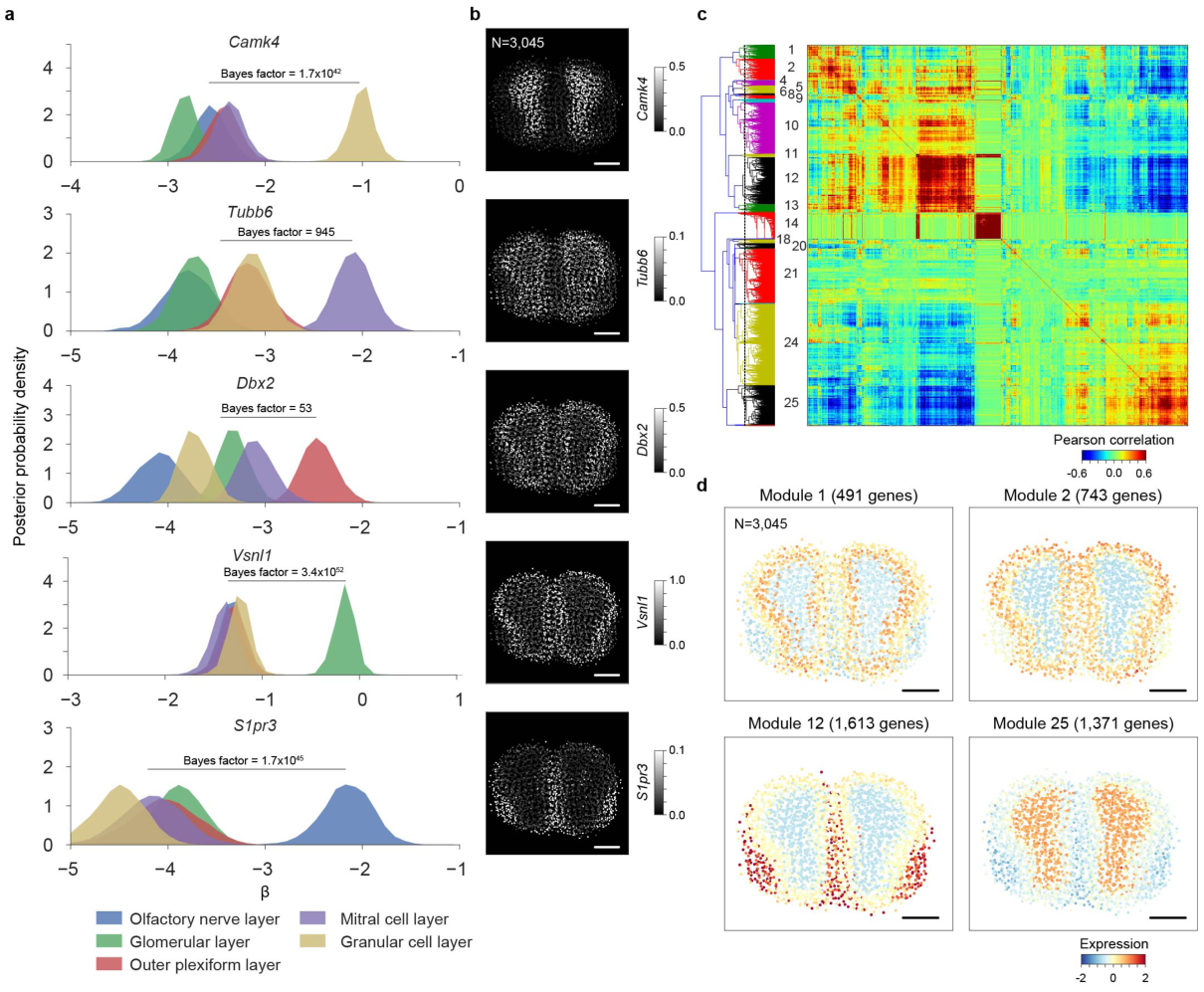
Detection of spatial gene expression patterns in mouse main olfactory bulb at gene and transcriptome levels. (**a**) The posterior distributions of the region-specific coefficient parameters β of *Camk4, Tubb6, Dbx2, Vsnl1*, and *S1pr3* in mouse main olfactory bulb. The color key is illustrated at bottom. The Bayes factors to quantify the difference between the right-most region and the combination of other regions are listed. (**b**) Spatial gene expression of *Camk4, Tubb6, Dbx2, Vsnl1*, and *S1pr3* in mouse main olfactory bulb are visualized using the estimates obtained using the integrative spatiotemporal approach. The number of ST spots is listed. Scale bar is 800 μm. (**c**) Biclustering of the mouse main olfactory bulb ST data reveals spatially co-expressed genes. The color visualizes the Pearson correlation coefficients between pairs of genes. The analysis considers 13,340 genes. The dashed vertical purple line in the dendrogram denotes the break. The identifiers given to the co-expression modules are listed on left. (**d**) Spatial dynamics of co-expression modules 1, 2, 12, and 25 containing 491, 743, 1,613, and 1,371 genes, respectively, in mouse main olfactory bulb are visualized. The colors represent averages (across genes) of standardized expressions (across spots within genes) at each spot. The number of ST spots is listed. Scale bar is 800 μm.

**Supplementary Figure 9.**
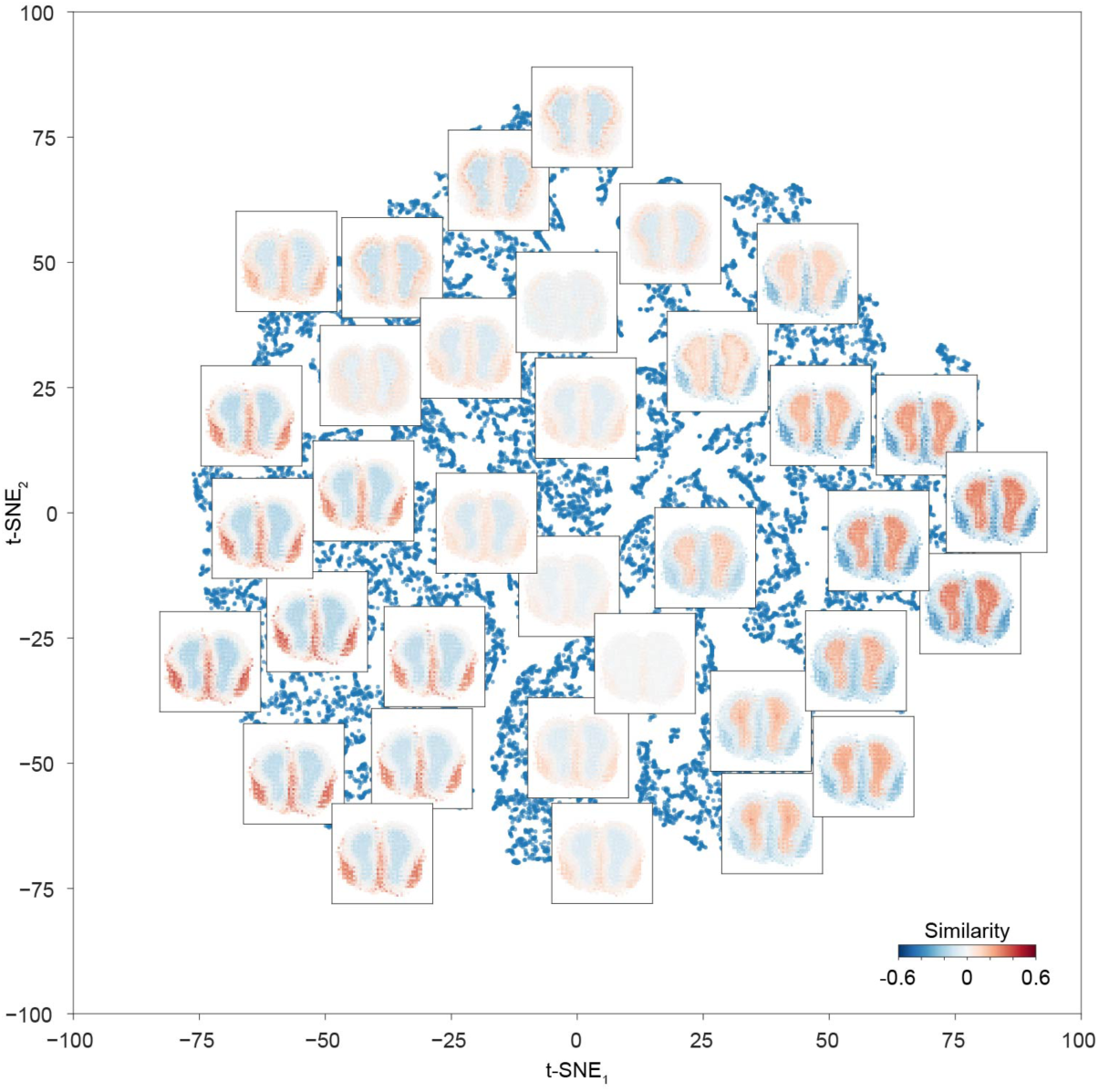
Localization of mouse main olfactory scRNA-seq data. A visualization of the localization maps of the mouse main olfactory bulb scRNA-seq data using t-SNE. Localization maps are visualized for selected cells. Each localization map is calculated based on 3,045 ST spots. Cells are not assigned cell type labels.

### Acknowledgements

The study was supported by Target ALS, The ALS Association (grant no. 15-LGCA-234), The Tow Foundation, the Knut and Alice Wallenberg Foundation, and the Simons Foundation. S.V. is supported as a Wallenberg Fellow at the Broad Institute of MIT and Harvard. We thank the Scientific Computing Core of the Flatiron Institute for computational resources.

## Contributions

T.Ä. and R.B. developed and implemented Splotch. S.M. and S.V. performed the experiments, with help from K.K., M.C., and C.B. T.Ä., S.M., and S.V. analyzed the data. All authors discussed the results and wrote the manuscript.

